# Reinforcement Learning via Brain Feedback for real-time fMRI-based adaptive stimulus generation

**DOI:** 10.64898/2026.08.08.743648

**Authors:** Giuseppe Gallitto, Robert Englert, Balint Kincses, Raviteja Kotikalapudi, Jialin Li, Kevin Hoffschlag, Sulin Ali, Ulrike Bingel, Tamas Spisak

## Abstract

Traditional fMRI studies rely on predefined task paradigms, where fixed stimulus designs limit the flexibility with which brain-stimulus relationships can be explored.

Here, we introduce Reinforcement Learning via Brain Feedback (RLBF), a framework and open-source software package for adaptive stimulus optimization using real-time fMRI. RLBF reverses the conventional direction of inference by using neural responses to guide the exploration of stimulus spaces through reinforcement learning, enabling optimization of predefined brain targets such as regional activity or multivariate neural signatures.

The accompanying Python-based software provides a modular framework integrating real-time fMRI data processing, reinforcement learning agents, adaptive stimulus generation, simulation-based testing, and experiment monitoring. Its flexible architecture allows researchers to customize preprocessing pipelines, reward functions, stimulus spaces, and RL strategies for diverse closed-loop neuroimaging applications. We validate the framework in a proof-of-concept study (N=10), demonstrating real-time optimization of a simple visual stimulus space by adapting checkerboard contrast and frequency to maximize primary visual cortex (V1) responses within a single 10-minute fMRI session.

RLBF provides an extensible foundation for brain-guided stimulus optimization and enables new approaches for investigating neural specificity, individualized brain–stimulus relationships, and adaptive experimental design.

## 1. Introduction

In traditional task-based functional magnetic resonance imaging (fMRI) studies, choosing the right experimental paradigm is key to identifying neural correlates of various psychological, cognitive or behavioral processes. The paradigm is essentially a formal statement about the research question, based on prior knowledge of the underlying processes, giving its construct validity [1]. However, using “hand-crafted” paradigms that are rooted in implicit assumptions imposes a significant limitation on the interpretability of the results [2]. These hidden assumptions mean that the interpretation of the identified neural correlates is constrained by the paradigm’s implicit theoretical and conceptual framework. For instance, observing activation in the amygdala during the fearful-face task does not imply that the amygdala specifically encodes fear, as the same region is recruited by many other psychological processes [3]. This is a manifestation of the reverse inference problem; the flawed practice of assuming that a specific brain region’s activation is directly and uniquely linked to a particular cognitive function [2,4], which is still a widespread issue in the neuroimaging literature and can only be partially overcome by multivariate encoding and decoding techniques [5].

Here, we present the concept, implementation, and initial proof-of-concept results of an alternative task-based framework for investigating brain–behavior relationships, in which the expression of a behavioral or cognitive process of interest is not restricted to a single predefined set of stimuli. Instead, our approach uses feedback signals derived from a closed-loop real-time fMRI setup to dynamically optimize stimulus parameters within a defined “stimulus space” to maximize an arbitrary target brain readout. Thereby, our approach - termed Reinforcement Learning via Brain Feedback (RLBF) - reverses the conventional direction of inference in task-based neuroimaging (from stimulus-to-brain toward brain-to-stimulus), providing a framework for adaptively identifying stimuli that optimally engage specific neural representations. This may enable more direct characterization of psychological or cognitive processes associated with a given brain feature while reducing the constraints imposed by hand-crafted experimental paradigms.

Our concept builds upon the pioneering “automatic neuroscientist” framework of Lorenz et al. [6], which introduced the idea of using real-time fMRI to adapt experimental paradigms online. We extend this principle into a general reinforcement learning framework [7], allowing adaptive optimization over arbitrary stimulus spaces rather than predefined task parameters. Leveraging RL’s ability to learn adaptive policies in dynamic environments, RLBF can be applied to complex stimulus spaces, including the latent spaces of generative models, thereby enabling the discovery of stimuli that maximize the specificity and interpretability of neural readouts.

In the present manuscript, we first provide a general formalization of the RLBF concept and introduce a streamlined open-source software implementation. We then illustrate the concept and validate the implementation in a proof-of-concept study focusing on the optimization of a deliberately simple two-parameter visual stimulus space. Finally, we discuss opportunities and limitations for extending the framework toward broader applications and increasingly complex stimulus spaces, enabling adaptive exploration of stimuli with rich and unconstrained characteristics.

## 2. Concept

As an alternative to assigning brain activity patterns to predefined behavioral or cognitive processes (formalized as sets of experimental stimuli or “paradigms”), the RLBF approach aims to adaptively identify the psychological or cognitive functions associated with a predefined brain feature (e.g., activity within a region, connectivity within a network, or a multivariate predictive readout). This is achieved through the integration of real-time fMRI (Figure 1, bottom) [8], reinforcement learning (RL; Figure 1, top left) [7], and flexible stimulus-space modelling (Figure 1, top right). Specifically, the reinforcement learner controls the parameters defining the stimulus space and presents the resulting stimuli to the participant. The predefined brain feature of interest is then estimated from real-time fMRI data and used as either the input signal or reward function for the RL model, thereby closing the adaptive loop. This enables the optimization of a broad range of stimulus spaces, from simple parametrized visual features to complex dynamic environments, or even the generation of entirely novel stimuli through latent spaces of generative models, with the goal of identifying stimuli that reliably elicit targeted neural responses.

**Figure 1:**
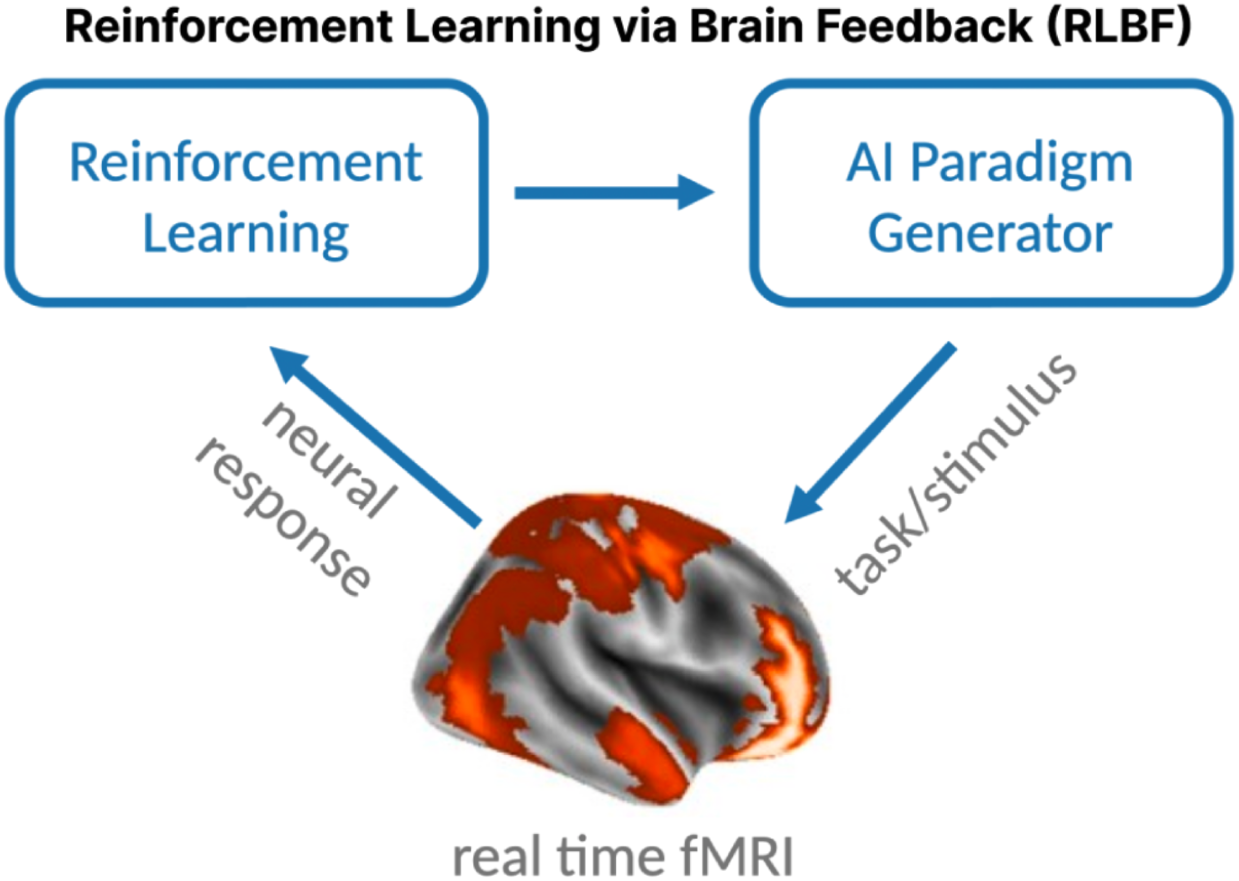
Concept behind RLBF. RLBF replaces fixed, hand-crafted experimental paradigms with an adaptive framework in which stimuli are represented as points within a continuously optimizable stimulus space. A reinforcement learning agent explores this space and updates stimulus parameters based on reward signals derived from target brain features estimated in real time. Through this closed-loop optimization, RLBF identifies stimulus configurations that maximize specific neural readouts, including regional activity, network connectivity, or multivariate predictive signatures.

Our approach builds on and integrates several methodological developments that use neural signals to identify, characterize, or optimize stimuli capable of modulating brain activity and probing brain function. It connects to classical closed-loop neuroimaging approaches [6] while also extending more recent efforts that leverage generative AI to identify stimulus-selective brain representations (e.g., [9,10]). By introducing closed-loop optimization into these frameworks, RLBF provides a principled approach for adaptively exploring stimulus spaces and identifying stimulus configurations that maximize targeted neural responses. Thus, RLBF promises to advance our understanding of the relationship between brain activity and stimulus features by enabling the discovery of novel and potentially more effective stimulus configurations that may not be captured by traditional, hand-crafted paradigms.

## 3. Real-time fMRI Software

We translated the general concept of RLBF into a modular and extendable software implementation [11] that (i) offers quick adaptability by being compatible with built-in real-time data export features of MRI scanners (currently compatible with Siemens scanners); (ii) provides a fast and lightweight real-time fMRI processing pipeline; (iii) favors transparency against complexity by implementing a simple yet robust and highly interpretable reinforcement learning approach (soft Q-learning) [12]; (iv) provides a flexible “container” for arbitrary stimulus space implementation (with a simple 2-parameter visual stimulus space as an example) and (v) supports real-time experiments with an interactive graphical dashboard.

While the current implementation is tailored for our proof-of-concept study (i.e. it uses a specific RL algorithm and a simple, two-parameter example stimulus space), the software architecture is intentionally built for modularity and extensibility. All these modules can be easily extended or even replaced to meet the needs of a wide variety of experiments. For instance, experimenters can effectively “plug in” a learning framework of choice without requiring modifications to the fMRI data-handling pipeline. This flexibility allows for the creation of RLBF experiments with varying levels of complexity, from simple stimulus optimization to more sophisticated cognitive control paradigms, while maintaining a consistent and reproducible experimental pipeline governed by the software’s core functionalities. Thereby, our software framework allows easy access to the RLBF technique and lays the groundwork both methodologically and practically for future studies, towards potentially more complex, AI-driven stimulus optimization applications.

### 3.1. Relation to existing real-time fMRI software

To date, existing software options for real-time fMRI remain limited in their flexibility and scope. As described in [13], many available solutions require commercial licenses, depend on specific software environments, or constrain analyses to predefined settings, such as focusing on a single region of interest. Moreover, most existing real-time fMRI frameworks have been developed primarily for neurofeedback applications, where minimizing data transmission and processing latency is essential: excessive delays between brain activity and feedback presentation can impair participants’ ability to learn the desired self-regulation strategy.

In contrast, RLBF represents a different use case with distinct methodological requirements. Here, the primary challenge is not achieving the lowest possible latency, as delays that would interfere with human neurofeedback learning are generally less critical for reinforcement learning-based optimization. Instead, the focus shifts toward providing a flexible and extensible framework capable of supporting a broad range of adaptive experiments. RLBF studies may involve diverse target signals, reinforcement learning agents, optimization strategies, and stimulus spaces, ranging from simple parametrized stimuli to complex environments such as generative AI-based stimulus models or real-time interactive gaming environments.

This flexibility also introduces additional challenges, as reinforcement learning algorithms typically involve a large number of tunable components and hyperparameters. Directly optimizing these parameters during real-time human experiments would be inefficient and costly. Therefore, the RLBF framework was designed to support extensive simulation-based development and testing, allowing researchers to explore and optimize RL configurations in silico before deployment in real-time fMRI experiments. Consequently, the software architecture prioritizes modularity, transparency, and well-defined interfaces, enabling researchers to adapt and extend the framework across a wide range of RL-based in-silico simulations and in-vivo neuroimaging applications rather than optimizing it for a single neurofeedback scenario.

Although open-source real-time fMRI solutions are available and support integration with different scanners and experimental setups, they are generally not designed around the requirements of RL-based adaptive experimentation. Our software addresses this gap by providing native support for integrating reinforcement learning components - including custom agents, environments, and optimization strategies - into real-time fMRI workflows. Rather than relying on a monolithic application structure, the framework is built around a modular architecture consisting of abstract classes defining standardized interfaces and reusable core components. This design allows users to inspect, modify, and extend the codebase, facilitating the implementation of diverse RLBF experiments and enabling rapid iteration between simulation and real-time applications. The RLBF software package is publicly available at [11].

### 3.2. Implementation details

The primary goal of our implementation is to provide a framework capable of training reinforcement learning (RL) algorithms on incoming real-time fMRI data from individual participants, enabling RL agents to adaptively identify stimulus characteristics that drive targeted brain responses. In addition, the framework was designed to support simulation-based development and testing of the complete real-time pipeline, providing a critical step for optimizing experimental configurations before deployment in human experiments.

The software architecture consists of three key components: (i) a **controller**, which manages incoming fMRI data streams; (ii) a **workflow**, which orchestrates communication between the modules responsible for fMRI preprocessing, reinforcement learning, stimulus presentation, and experiment logging; and (iii) a browser-based, customizable **dashboard** for real-time visualization of fMRI data, RL agent performance, and experimental parameters, enabling continuous monitoring and interaction during experiments. All user-defined parameters and hyperparameters are centralized within a single configuration file, facilitating transparent setup, reproducibility, and systematic exploration of different experimental configurations.

The RLBF package was developed entirely in Python 3.11.7. The currently implemented preprocessing pipeline relies on established neuroimaging tools, including ANTs (Advanced Normalization Tools) [14], FSL [15], and dcm2niix [16]. Schematics illustrating the interaction between the framework components and a preview of the real-time dashboard are provided in Figure 2 and Supplementary Figure 5.

**Figure 2:**
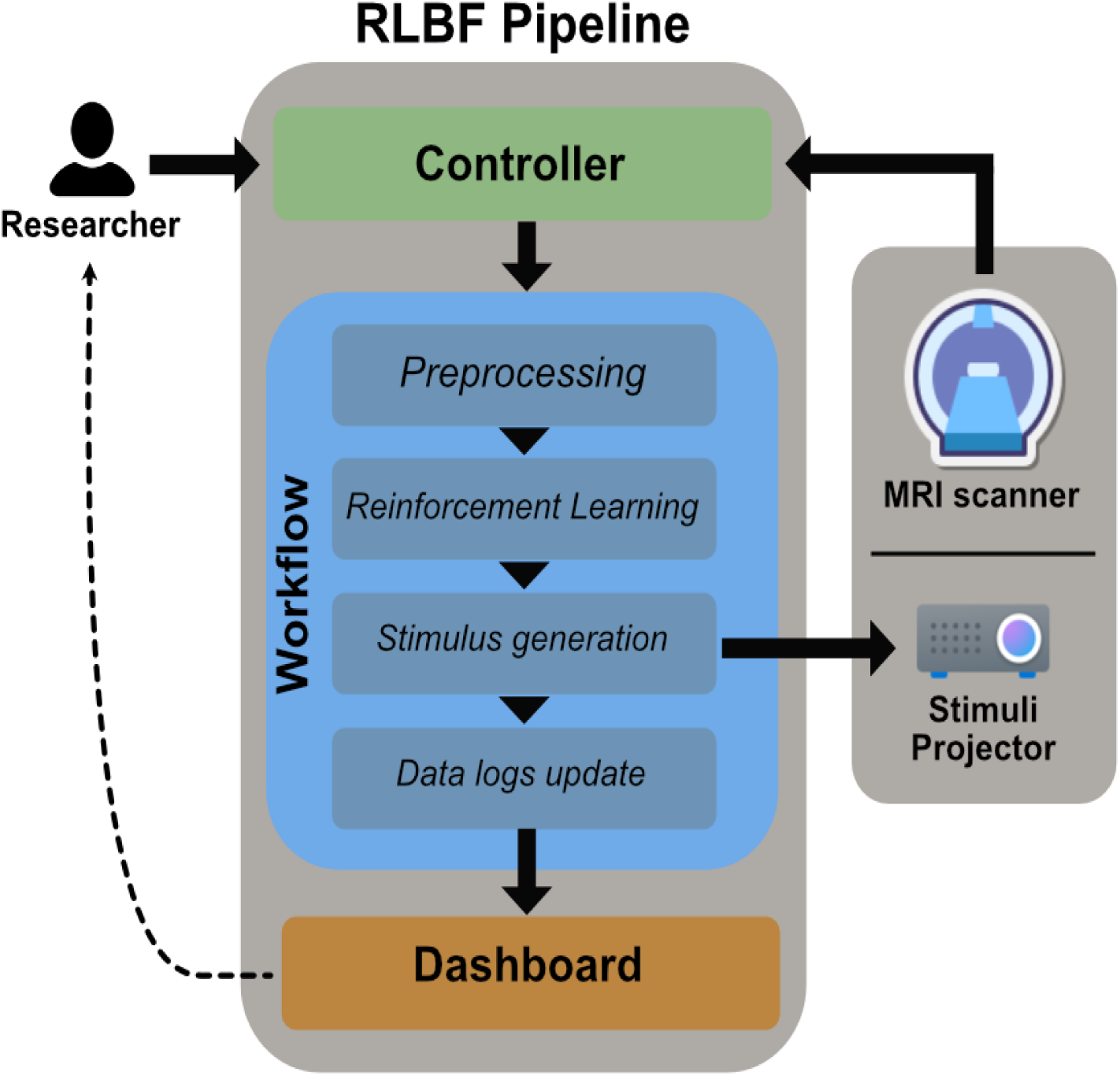
Schematic of the RLBF software’s [11] core components. The Controller component monitors incoming data streams and forwards newly acquired volumes to the Workflow component, which orchestrates communication between modules responsible for preprocessing, reinforcement learning, adaptive stimulus generation, and experiment logging. Each incoming volume is processed according to the selected preprocessing pipeline and stored for subsequent analysis. Once all volumes associated with a trial have been acquired, the resulting brain features are provided to the RL agent, which selects an action that is transmitted to the stimulus generation module (RL environment) to render the next stimulus. All relevant user-defined workflow data are continuously passed to the data logger, where they are stored on disk and made available for real-time visualization through the Dashboard component.

### 3.3. Controller Component

The Controller component manages the information flow at the highest level. Its only function includes orchestrating the handling of incoming fMRI volume data and sending it to the workflow component for real-time processing. It directly manages data coming from the fMRI scanner, receiving raw volume data as it is acquired. Currently the software supports the ‘indirect’ export of single DICOM volume data files for each volume separately to a user-defined network share using the underlying server message block (SMB) network protocol [17]. This is provided for a wide variety of MRI scanners by the manufacturer without further tools needed, i.e. it does not require running custom scripts on the console workstation. The present implementation was extensively tested on a Siemens MAGNETOM Vida 3T system. Note that while direct TCP/IP-based data transfer is faster and more consistent than indirect file-based export [17], we found that the speed and reliability of the indirect method is sufficient for the RLBF approach. Thus, the controller module’s data interface can rely on indirect export, making it not only compatible out-of-the-box with a wide range of scanners, but also significantly lowering the barrier for researchers who have limited experience with real-time fMRI to perform RLBF.

### 3.4. Workflow Component

The Workflow component is the computational engine for the real-time fMRI closed loop. It receives each individual volume from the Controller Module and manages the data flow between specialized **workflow sub-components**, which implement specific units of the real-time fMRI pipeline.

These four specialized subcomponents are:

1. **Preprocessing pipeline**: handles fMRI data preprocessing at any stage, including before (e.g., reference or localizer volumes) and during real-time data acquisition. The preprocessing pipeline is also tasked with the in-memory storage of temporary derivatives (e.g. trial-related fMRI and motion data).
2. **RL Agent**: implements a specific Reinforcement Learning (RL) agent. Its function is to select parameters that determine the characteristics of the next stimulus (**action**) to be presented in the scanner and receive the preprocessed fMRI data as input or to calculate a reward signal and update its internal knowledge of the stimulus space.
3. **Stimulus generation (RL environment)**: receives stimulus parameters from an agent and renders the corresponding stimulus for presentation to a human participant. If Generative AI is used, this sub-component is also tasked with containing the inference process for stimulus generation (e.g. through an API).
4. **Data logs**: capture all information generated during program initialization and real-time acquisition, logging it to a file on the disk. The log file serves both as a real-time data source for the Dashboard core component and as a comprehensive record of the experiment. It stores relevant information about stimulus parameters, RL agent states, brain-derived signals, experimental events and system parameters, enabling subsequent analysis, reproducibility, and detailed bookkeeping.

#### 3.4.1. Workflow: Preprocessing Pipeline

Upon program initialization, the pipeline sub-component prompts the user to select a specific preprocessing pipeline. The current implementation (named “*standard pipeline*”) offers decisions such as the type of image registration to the standard template (defaulting to symmetric diffeomorphic image registration - SyN [18]) and the method for selecting ROIs. ROIs can be selected from an atlas or drawn as spheres from user-specified coordinates in native-space, a particularly useful functionality in cases where the registration to a standard space might be challenging or when custom regions are desired in the absence of an atlas or during pilot studies.

The main operational steps of the standard pipeline are as follows: it first establishes reference volumes – either a single functional volume or a combination of structural and functional volumes. The functional reference volume, or the structural one when available, is registered to the standard template, to generate a transformation matrix. The importance of this matrix is twofold. First, it allows the alignment of all subsequently acquired functional volumes to the structural template, a straightforward option when using simple linear registration. Second, its inverse can be used to bring specific ROI masks into a participant’s native space, which is a better option when a non-linear registration to the standard space is chosen.

After the initial preprocessing of the reference volume(s), the derived processing parameters, the volumes themselves, and the transformation matrix to the standard template are all saved. This allows for their efficient reutilization in successive runs, saving time and computational resources. However, to maintain flexibility and accommodate potential changes in experimental design or subject positioning, experimenters will always be prompted to decide whether they wish to acquire new reference data and repeat the preprocessing steps each time the program restarts. After the preprocessing steps and the reference volumes are fixed, the pipeline sub-component focuses on data quality and precise alignment. An initial image alignment step is crucial, regardless of whether experimenters transform the ROI mask(s) to the participant’s native space or apply the transformation matrix to move functional volumes into standard space. This alignment employs the SyN method [18] as a default, preceding the subsequent motion correction phase.

Motion correction is performed using FSL’s mcFLIRT [19] to minimize the impact of head movement on the data by performing a fast registration of the last incoming volume to the functional reference. An important byproduct of this step is the calculation of motion regressors, which provide a quantitative measure of head movement. These regressors are highly valuable, particularly in contexts where a General Linear Model (GLM) is used to estimate brain-based signals from a continuous block or sliding-window of data for the reinforcement learner, as they help accounting for variance related to motion. The pipeline sub-component also allows for the exclusion of brain volumes if the motion exceeds user-defined motion thresholds (defaulting to 0.25 mm motion for ≥50% of volumes in a trial), thereby preventing motion-derived noise from propagating to the reinforcement learner and the stimulus space.

Following alignment and motion correction, the single volume data is rank-harmonized to mitigate global signal effects, and the data to be used in reward calculation is then isolated using the user-defined binary brain mask and stored in memory. If the experimenter defined a trial as encompassing multiple volumes, the masked data is flattened and stacked into a 2D matrix (volumes × voxels). This format is particularly useful when reward calculation is

based on classical linear models. Alternatively, if a trial corresponds to a single volume acquisition, the flattened data is used directly to calculate the reward signal for the reinforcement learning algorithm. All the data received by the pipeline is converted to Nifti file format, before preprocessing, using dcm2niix [16], which ensures that each volume has proper orientation and affine matrix before image registration.

#### 3.4.2. Workflow: Reinforcement Learning

When preprocessed data are returned by the preprocessing pipeline, the Workflow component sends them to the RL agent sub-component, which is trained each trial (corresponding to each block in a block-design experiment or 1-to-N sliding-window volumes in event-related designs, depending on the experimental paradigm). The current implementation features a Soft-Q-learning algorithm [12] as the reinforcement learner. In Soft-Q-learning, the q-values are convolved with a gaussian kernel to model dependencies across similar actions, improving generalization and avoiding overfitting. Actions are then selected based on Softmax probability, so that higher Q-values are more likely to be chosen but lower Q-values still have a non-zero probability, promoting exploration. As the Q-values in the table converge towards the optimum during training, the algorithm naturally begins to exploit these higher values, focusing more on actions with higher expected rewards (exploitation). The Soft-Q-learner was chosen due to its simplicity and interpretability, as the Q-table can provide an instant, straightforward visual illustration of the learning process.

Upon the completion of data collection for a trial and the overall degree of head motion is evaluated, the pipeline sub-component determines if the data are suitable for the reinforcement learning algorithm. If motion exceeds the user-defined thresholds, the RL agent is instructed to discard the data, no reward is computed and the agent selects a new action based on its prior state. Conversely, if motion levels are acceptable, the reward for that trial is calculated and the internal state of the agent is updated. In the current implementation the reward is calculated by means of an Ordinary Least Squares (OLS) fitting the time series data of each trial (in a n-volumes block-design fashion) with the hypothesis function, derived by convolving the block’s timing (boxcar function) with a canonical hemodynamic response function (HRF) chosen by the experimenters. The model yields coefficients representing the activation amplitude (β_1_ , in arbitrary fMRI units) and an estimation of the baseline (β_0_ , the intercept), which allow to obtain a reward value (*r*) as follows:

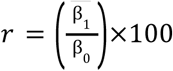

where β_1_ is the mean of the coefficients and *r* is a value analogous to percent signal change (PCS), which is empirically ranged approximately between 0 and 5. Based on this expected value range, the Q-table values are initialized, by default, to a value of 2.0 (around mid-range) before each real-time fMRI run. It is important to notice that this range can change due to image harmonization for intensity correction and the Q-table initialization value should be consequently adjusted to fit the new values.

For experiments in which the stimulus-response relationship does not follow a predefined block or event-related structure, an alternative reward estimation strategy can be implemented. Instead of fitting a fixed block-design model, the framework can be easily extended to use sliding-window approaches combined with hemodynamic response function (HRF) deconvolution to align continuous brain readouts with the precise timing of generated stimulus events. This enables the use of continuously varying neural features as reward signals, such as multivariate predictive signatures (e.g., the Neurologic Pain Signature, NPS), which can be dynamically matched to continuously modulated stimuli (e.g., visual environments or other parameterized stimulus streams). Such approaches further extend RLBF beyond traditional experimental paradigms by allowing optimization based on temporally resolved neural responses without requiring predefined trial structures.

To enhance signal-to-noise ratio, the current implementation of the reward calculation allows the selection of the voxels with the highest *K* coefficients to be utilized in reward computations. This approach can be situationally beneficial in a simple RL framework, where we retrain the agent for each participant and run and do not require generalization across them. Consequently, what would be normally regarded as a selection bias, allows an improvement of the overall performance.

After a reward is calculated, our default implementation uses soft-Q learning to learn actions that maximize it. With this approach a so-called Q-table is updated to incorporate the new information. The Q-table update is performed at the position of the last action taken using the following equation:

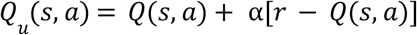

where Q(s,a) represents the previous Q-value (i.e., the expected reward) for the current state-action pair, α is the learning rate, and the term [*r* − *Q*(*s*, *a*)] corresponds to the temporal difference error [20], which quantifies the discrepancy between the observed reward and the previously estimated value and is used to update the Q-value.

To account for the continuous nature of many stimulus spaces and to promote generalization across neighboring actions, the Q-value update is not restricted to a single state-action pair. Instead, the update is spatially smoothed by convolving the Q-table with a Gaussian kernel, where the full width at half maximum (FWHM) of the kernel is a user-defined hyperparameter controlling the degree of generalization between neighboring actions. This allows information obtained from a single stimulus configuration to influence similar configurations while preserving the overall structure of the learned reward landscape.

Action selection is performed using a soft-Q learning strategy, in which actions are sampled according to a Boltzmann (softmax) distribution over the current Q-values rather than selecting only the action with the highest estimated value. Specifically, the probability of selecting action *a* in state *s* is given by:

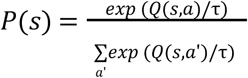

where τ is a temperature parameter controlling the exploration-exploitation trade-off. Higher values of τ result in more exploratory behavior by increasing the probability of selecting lower-valued actions, whereas lower values increasingly favor actions with the highest expected reward.

This stochastic action-selection strategy enables continuous exploration of the stimulus space while progressively focusing on configurations associated with stronger brain responses. Although the temperature parameter τ determines the general balance between exploration and exploitation, the effective exploration behavior is also shaped by the amount of evidence accumulated in the Q-table. As the agent acquires sufficient evidence and a clear high-value region emerges, the softmax distribution increasingly concentrates around the corresponding Q-value peak, resulting in a potentially near-deterministic action selection even at relatively high temperature values. Thus, the approach provides an adaptive exploration-exploitation trade-off that naturally evolves with evidence accumulation, allowing broad exploration during early learning stages and increasingly focused optimization as the reward landscape becomes better characterized.

A schematic diagram depicting reward calculation and Q-table updating in the current implementation is available in Supplementary Fig. 1. In future implementations, state-of-the-art reinforcement learning architectures could be employed either to optimize the characteristics of more complex, dynamic environments or to integrate with generative AI models. In the latter case, the models’ latent spaces could serve as the stimulus space, a strategy commonly observed in modern generative AI approaches, such as fine-tuning large language models [21] or image transformations [22,23].

#### 3.4.3. Workflow: Stimulus Generation

The RL agent output, i.e. the action defining the characteristics of the stimulus to be generated, are sent to the stimulus generator, i.e. the RL environment sub-component. The stimulus generation module is designed to be highly customizable and is expected to be substantially adapted or reimplemented by experimenters, as the definition of the stimulus space represents a central component of RLBF experiments.

In the current implementation, we provide a deliberately simple example consisting of a flickering checkerboard parameterized by contrast and frequency. This example serves both as a demonstration of the module’s functionality and as the stimulus space used in the proof-of-concept study presented later in the present manuscript. The two parameters are defined with a range of values between 0 and 1.0, where a contrast of 0 means that the checkerboard is not visible, while a contrast of 1.0 corresponds to the brightest stimulation possible. On the other hand, a frequency of 0 means that the checkerboard does not flicker, while a frequency of 1.0 corresponds to the maximum flicker rate (close to being perceived as a static image again, with attenuated contrast). The maximum flicker rate is defined, by default, as a reference value (Rf) of 30Hz and the actual flicker rate (F) for any frequency value f between 0 and 1.0 is calculated using the following equation: *F* = *f*×*R_f_*.

Regarding contrast levels rendered by the stimulus generator, these are spaced non-linearly to account for the logarithmic relationship between true and perceived stimulus contrast and the corresponding neural responses, including potential saturation effects at higher contrast levels [24,25] (for details and the code about contrast/frequency scale calculation see Supplementary Fig.2 and our supporting repository [24]). The stimulus generator is represented by a custom RL environment developed in Pygame 2.6.1 [26].

#### 3.4.4. Workflow: Data Logs

The RLBF framework incorporates a data logging system, designed to easily record pertinent experimental data and variables available to the Workflow component and its sub-components. This system provides information regarding the reinforcement learning agent’s last and current actions, reflecting the agent’s exploration of the stimulus space, along with reward metrics, RL model convergence status, and motion regressors (rotations and translations). The logging sub-component allows the monitoring of the stimulus parameters (contrast and frequency, in our case), and facilitates the generation of visual reports through the Dashboard component. The main purpose of the logging functionality is to provide access to all relevant variables used during real-time scanning, allowing experimenters to record detailed data pertaining to the experimental paradigm and the real-time preprocessing pipeline for comprehensive visualization. The default logging sub-component is designed to access all data from the pipeline and the agent available for logging whenever a trial concludes and the Reinforcement Learner’s knowledge is updated. Specifically, all information of interest, generated by the real-time processing, is structured in a single JSON file that serves as the primary data source that is read by the Dashboard component for immediate visualization and monitoring of the ongoing fMRI acquisition and RL training.

The workflow system is designed for high efficiency, multi-threaded processing and has been tested with a 1-second Repetition Time (TR), enabling truly rapid feedback loops. Achieving this performance threshold doesn’t exclude the possibility of reaching faster processing times. However, these may vary depending on the complexity of the implemented workflow, the hardware at hand and specific computational demands.

### 3.5. Dashboard Component

The Real-Time Dashboard provides an intuitive and highly customizable web-based interface for visualizing the progress of the fMRI acquisition and the ongoing reinforcement learning training. Its core function is reading the output JSON file, generated by the Workflow component, and allowing users to monitor key metrics and feedback in real-time. A key strength of the dashboard lies in its flexibility, enabling users to interactively plot data using widely used Python packages such as Plotly [27] and Matplotlib [28]. This allows for the creation of rich and dynamic visualizations, ranging from visualizing time-series of BOLD activity to statistical summaries of RL-related and motion parameters. Beyond graphical representations, the dashboard also facilitates the display of tabular data (e.g., pandas DataFrames [29]) and printing important information via Markdown. The default implementation of the dashboard incorporates re-usable features for temporal navigation, allowing the exploration of progress in time by moving between old and newly acquired trials. The Dashboard component is built in Streamlit [30] with responsiveness in mind and empowers users to define a custom layout, allowing them to arrange plots, data tables, and text elements according to their specific monitoring needs and preferences. This level of customization ensures that the dashboard can be tailored to a wide array of experimental designs and requirements. A graphical representation of our specific Dashboard setup is illustrated in Supplementary Figure 5.

### 3.6. Compatibility with RL agents and environments

A deliberate architectural choice integrates the Reinforcement Learning (RL) and the stimulus generation sub-components within the workflow component. Specifically, both the sub-components can be fully user-defined, allowing experimenters to implement and integrate their own learning pipeline of choice with other functionalities of the real-time processing. This modularity ensures that the software is not constrained to a particular problem domain, fixed set of RL algorithms, or specific packages that implement standard reinforcement learning procedures (e.g. Models compatible with specific machine learning frameworks, Gymnasium environments [31], etc), and simultaneously enables experimenters to select any of these tools with minimal, if any, necessary modifications. Nonetheless, the software includes the Soft-Q learning agent [12] and the flickering checkerboard RL environment (stimulus generator), which act not only as re-usable tools but also as examples of how those sub-components should be implemented. All sub-components were specifically developed and utilized for our proof-of-concept study (see the “Proof-of-Concept Study” section).

### 3.7. Future directions regarding software implementation

The software is tested and validated using real-time data export functionalities available on Siemens scanners (i.e., Magnetom Vida 3T). However, although supporting the ‘indirect’ export of fMRI data [17] provides a useful initial pathway toward compatibility with multiple MRI platforms, we acknowledge the need for broader scanner support. Future development efforts will focus on three key areas: (i) expanding support for official integration of data streams from other major scanner providers, (ii) implementing direct export capabilities, and (iii) incorporating more advanced online processing techniques. Additionally, we will develop and evaluate new RL agents and environments to further enhance the robustness of the RLBF pipeline and improve its adaptability and generalization across diverse experimental scenarios.

By open-sourcing the software, we aim to enable and encourage community-driven development that can accelerate these efforts beyond the scope of a single development team. Such collaborative contributions may facilitate adaptation to a broader range of scanner platforms and lead to the creation of a growing ecosystem of reusable modules, including reinforcement learning agents, stimulus-space definitions, preprocessing pipelines, dashboard components, and reward functions. This modular and community-driven approach will be essential for supporting the diverse experimental requirements of future RLBF applications.

While still in early development, the software already provides a fully functional platform integrating online fMRI data processing with reinforcement learning, enabling direct access to online stimulus optimization for studies requiring simple stimulus generation and minimal fMRI preprocessing out of the box.

## 4. Proof-Of-Concept Study

The presented implementation of the software framework was validated in a simple proof-of-concept study, where we use real-time blood-oxygen-level-dependent (BOLD) response measurements from the primary visual cortex (V1) to provide feedback for the Soft-Q-learning algorithm [12], which is tasked with optimizing the contrast and frequency of a flickering checkerboard to maximize neural activity in V1. The algorithm iteratively adjusts the stimulus parameters to achieve the maximal BOLD response. While our framework was designed to implement arbitrarily complex stimulus spaces, in this proof-of-concept study, we deliberately utilize a very simple stimulus space where we have access to a reliable prior knowledge about optimal parameter values. Specifically, as the activity in V1 is known to be characterized by stable visual contrast and frequency sensitivity functions [24,32,33], we let the reinforcement learner manipulate the contrast and frequency of the stimulation. Building on the well-characterized responses of the primary visual cortex (V1) to checkerboard stimuli, our primary goal in this proof-of-concept study is to demonstrate the feasibility and effectiveness of the RLBF framework in a simple task and to validate the accompanying software implementation. Specifically, we aim to demonstrate that the concept at the base of RLBF, combining real-time fMRI and reinforcement learning, can be successfully integrated to optimize stimulus parameters. Furthermore, we assess the potential for deploying this approach at the level of single participants.

### 4.1. In-silico parameter optimization

Before bringing the checkerboard task to the scanner, we initially conducted a hyperparameter optimization process using both computer simulations and empirical data. For the simulations, we established a ground truth reward landscape to serve as a known target for the RL agent. This landscape was defined such that the reward increased linearly with contrast and non-linearly with frequency, peaking at the arbitrary frequency parameter value of 0.7 (see Figure 3a; further details in Supplementary Fig.2a). This specific ground truth was used solely within the simulation environment to generate the reward.

**Figure 3:**
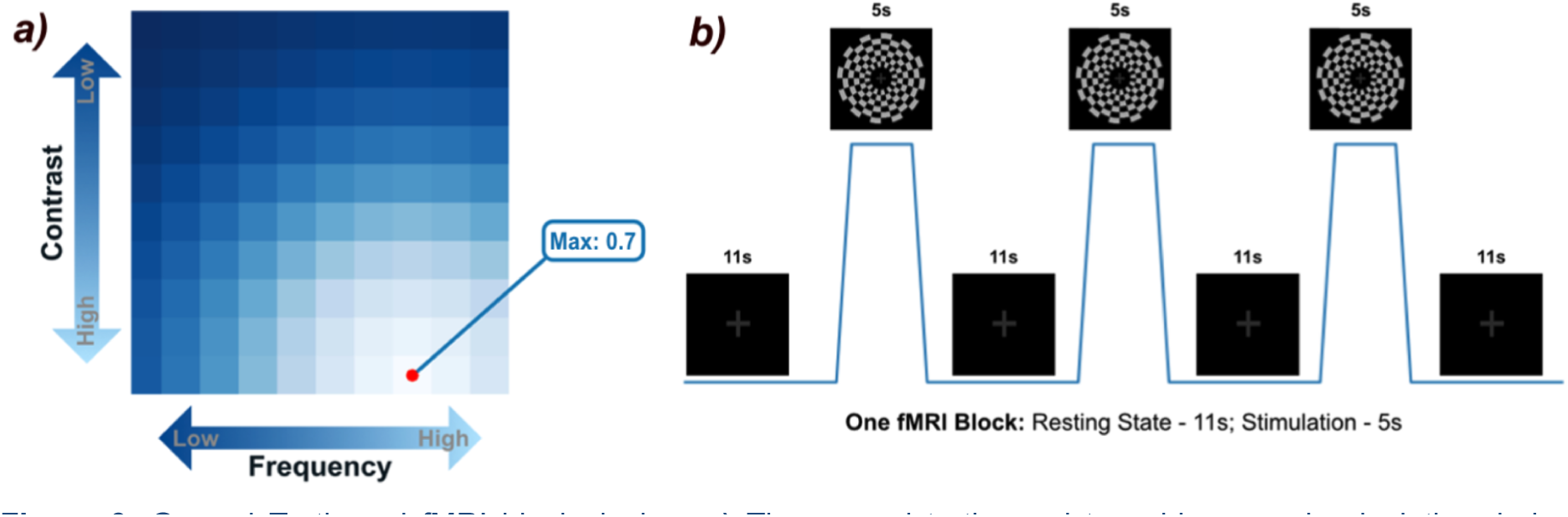
Ground Truth and fMRI block design. **a)** The ground truth used to guide reward calculation during computer simulations. As the reinforcement learner explores the paradigm space, it receives higher rewards when its actions align with or are near the optimum (high contrast and a frequency parameter of 0.7, shown as the maximum value in the table); **b)** the block design used during real-time fMRI acquisitions with human participants. In each block (resting state + stimulation), the reinforcement learner selects the contrast and frequency, and this process repeats for 35–45 blocks, corresponding to approximately 10 minutes of scanning time.

To determine the optimal hyperparameters for the RL agent, we conducted computer simulations. Each simulation run involved the reinforcement learner model completing 100 trials. In each trial, the agent selected a contrast/frequency combination and received a reward based on proximity to the ground truth’s optima (contrast: 1.0; frequency: 0.7), with added Gaussian noise to mimic experimental conditions. We performed multiple simulation runs using a grid search algorithm to optimize key hyperparameters influencing the reinforcement learner’s performance. This involved evaluating different configurations for an artificial signal-to-noise parameter (SNR), the Q-table smoothing parameter σ (which aids generalization across similar actions), and the learning rate α (which controls the speed of Q-value updates). To reduce the search complexity, other parameters like the Softmax temperature τ (which balances the exploration-exploitation trade-off in action selection) were kept constant during this simulation phase. The goal was to identify the hyperparameter set that allowed the Soft-Q-learner to properly estimate expected future rewards based on the simulated feedback.

The simulation-based hyperparameter search identified the following initial optimal values: a Q-table smoothing σ of 4.0 and learning rate α of 0.05. The fixed Softmax temperature τ was set to a value of 0.2. Fine-tuning the SNR showed that a parameter value of 2.0 or higher would result in the best Q-value increase at the position of the ground truth optima. For this reason, we fixed the SNR parameter to 3.30, a value that was selected based on approximate noise levels observed in preliminary empirical data, estimated post-hoc using the reward obtained during real-time fMRI runs. Calculating an SNR proxy measure from the reward was possible because the reward itself is derived directly from the fMRI data, and reflects the data’s inherent noise. Estimating the true SNR from fMRI data is non-trivial and our estimation method presents limitations, however our SNR proxy measure allows us to approximate the influence of lower or higher noise conditions within the simulation environment. The empirical SNR guiding the simulation value was calculated for each run using the following equation:

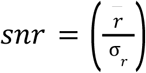

where *r* is the mean reward obtained in a participant’s run and σ*_r_* is its standard deviation. For more details about hyperparameter tuning and the reward ground truth, see Fig. 3a and Supplementary Fig. 2 and 3; detailed results are available in our supporting repository [34].

### 4.2. In-vivo fine-tuning

We further refined the hyperparameters using empirical data. This step is crucial to ensure that the hyperparameters are not sub-optimal when applied to real data and to optimize our real-time fMRI methodology.

After testing the hyperparameters obtained through the computer simulations on empirical data, we decreased the learning rate to 0.02, to reduce the algorithm’s sensitivity to the noisy neural feedback, and the temperature to 0.08 to encourage more exploitation and faster convergence towards an optimum in the stimulus space. These adjustments still allow the learner to sample a broader range of stimulus combinations before reaching an approximate optimal solution. We then tested these adjusted hyperparameters by re-running the simulations to verify their effectiveness (see Figure 4a).

**Figure 4:**
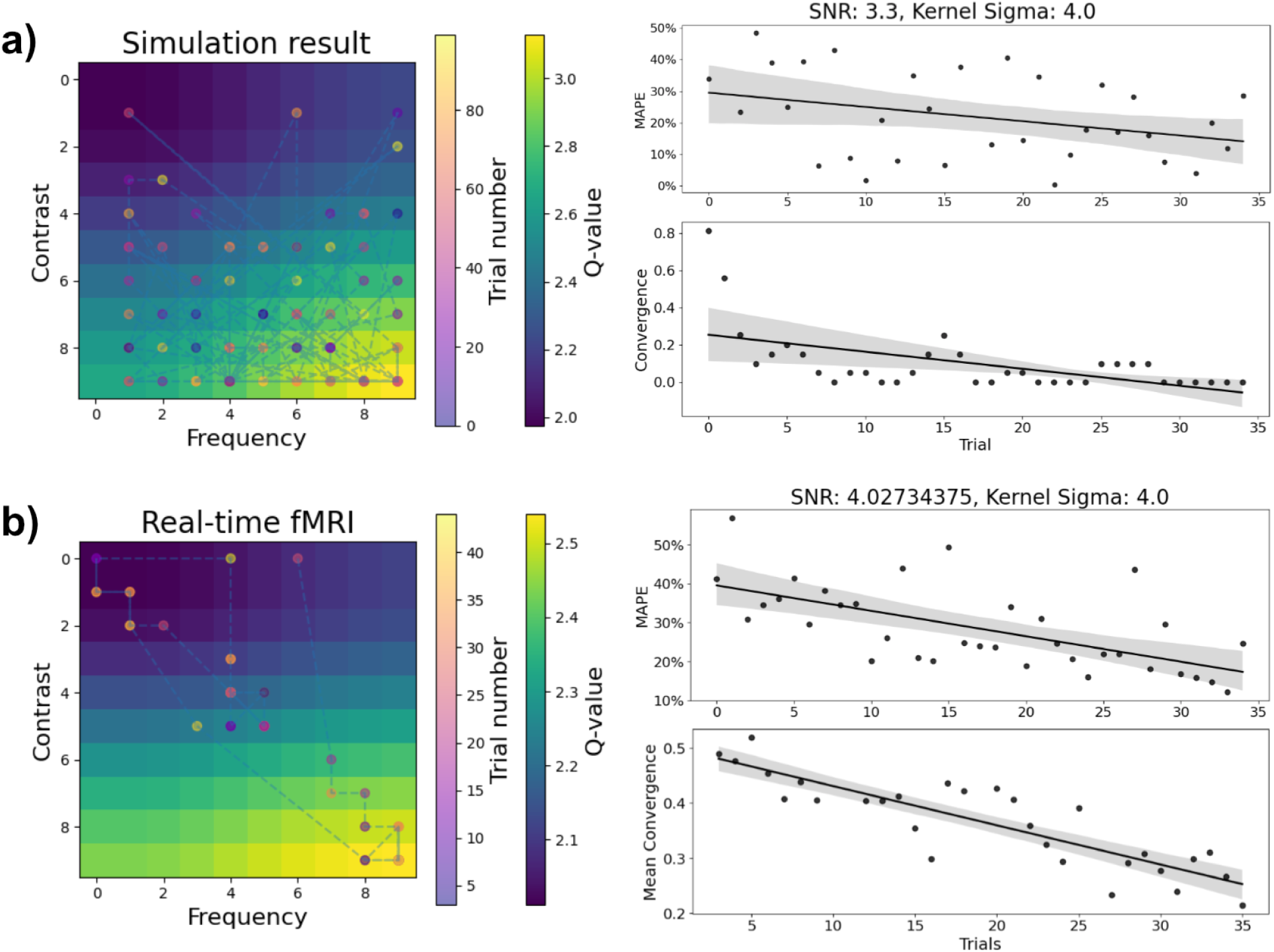
Optimization of the stimulus space by the RL agent. The figure shows the final Q-table obtained after 100 trials in the simulation (**a**) and the average Q-table across all participants (N=10) who underwent fMRI scanning (**b**). The Q-tables illustrate the trajectory of the reinforcement learner through the stimulus space as it optimizes contrast and frequency to maximize brain activity in V1. Mean absolute percentage error (MAPE) and convergence analyses demonstrate that the algorithm progressively reduces prediction error and rapidly converges toward the optimal region of the stimulus space (i.e., high contrast), reaching convergence after approximately 35 blocks (trials) on average. Shaded areas in the MAPE and convergence plots indicate the 95% confidence intervals of the regression estimates (y = values, x = trials). Although convergence patterns are highly consistent across simulation runs and participants, prediction error remains variable because the algorithm continues to explore the stimulus space throughout the experiment. This residual exploration is reflected in the color-coded trial progression and MAPE trajectories, and is expected given the stochastic action-selection strategy of the RL agent. Importantly, only a small proportion of trials exceeded the predefined motion threshold and were discarded, allowing the agent to continuously update its estimates. Code for interactive simulation and detailed analyses of individual participant trajectories and convergence behavior are available in the supporting repository [34].

### 4.3. Real-time data processing protocol

For our simple use-case, we selected the single functional reference preprocessing from the ‘standard pipeline’ implemented in the software. We generated the transformation matrix through linear registration to the standard MNI-152 template (2mm resolution), which was then applied to each subsequently acquired functional volume, enabling alignment to standard space. Finally, motion correction was performed on the aligned volume. For this proof-of-concept, participants were positioned with padding inside the scanner to minimize head motion; consequently, we retained all acquired volumes without having to apply the software’s motion threshold functionality. To isolate data of interest, we utilized the primary visual cortex (V1) ROI defined anatomically using the Brodmann atlas provided within the MRIcron software [35].

As a reinforcement learning agent, we employed our Soft-Q learning algorithm, incorporating a Gaussian kernel to model action dependencies and employing Softmax probability selection to balance exploration and exploitation, as previously described in the implementation section. By running the OLS model at the end of each trial on the acquired data, for reward calculation we utilized the voxels corresponding to *K* = 10% best coefficients in order to have higher SNR and speed-up training times. The Q-table, alongside key parameters (contrast, frequency, reward, convergence, and motion levels), were used to monitor data quality, track the learning process, and monitor the algorithm’s exploration of the stimulus space (See Supplementary Figure 5 for a screenshot of the dashboard as it was set-up for the proof-of-concept’s real-time data acquisition).

Regarding the stimulus generation environment, utilized during real-time fMRI acquisition, participants were presented the flickering checkerboard with parameters (i.e., contrast and frequency) controlled by the reinforcement learning agent. The agent dynamically adjusted these parameters at the end of each trial to optimize learning, expecting that it would identify high contrast as the primary characteristic driving maximal activity within V1 while simultaneously adjusting frequency to reflect individual sensitivity to temporal frequencies and maintain the overall activity high.

### 4.4. Experimental design

The experimental design for real-time fMRI acquisition with participants encompasses multiple trials, each performing a full loop in the proposed system (Figure 1). Each trial consists of resting (11 seconds) and stimulation (5 seconds) blocks. The resting condition involves a fixation cross at the center of the screen, while the stimulation condition features both the fixation cross and the flickering checkerboard (see Figure 3b). The presentation happened via an MRI-compatible projector (resolution 1920x1080 pixels) projecting onto a screen positioned at the rear of the scanner bore. Participants viewed the screen via a mirror mounted on the head coil, with an approximate viewing distance of 120 cm. The circular checkerboard stimulus had a diameter of approximately 30 cm, covering a visual angle of 14.25°. A central fixation cross (4 cm diameter) was overlaid on the checkerboard during stimulation blocks. We confirmed with participants that they could comfortably view the entire checkerboard stimulus and that the stimulus would fill most of their field of view while maintaining gaze on the fixation cross, which they were instructed to do throughout the experiment.

### 4.5. MRI data acquisition

We acquired fMRI data on a Siemens Magnetom Vida 3T scanner using an Echo Planar Imaging (EPI) sequence accelerated with Simultaneous Multi-Slice [36] (SMS; TR=1s; 3mm slice thickness; 76x76x45 matrix). Real-time runs typically involved 35-45 blocks, lasting approximately 10 minutes. During these runs, the RLBF framework’s logging sub-component recorded all data processed by the online analyses. All log data (for each participant and run) is publicly available in our supporting repository [34], which includes a dashboard to visualize trial-by-trial training progress.

### 4.6. Study participants

A total of N=10 young healthy volunteers (M=5, F=5, mean age=25.9, age SD=3.93) were involved in the proof-of-concept study. All participants underwent thorough screening as epilepsy, metal implant, non removable piercings, pacemaker, tattoo in head/neck position, pregnancy or claustrophobia were considered as contraindications for MR measurement. The study was conducted in accordance with the Declaration of Helsinki, complies with all relevant ethical regulations for work with human participants and has been approved by the local or national ethical committees. All participants gave written informed consent before testing. N=3 participants were engaged in the in-vivo hyperparameter optimization process to refine the results from the in-silico hyperparameter optimization (see the “In-vivo fine-tuning” section for details). The remaining N=7 volunteers were scanned exclusively using the resulting optimal hyperparameters.

## 5. Results

We primarily assessed the outcomes of our proof-of-concept study using three metrics: (i) the mean absolute percentage error (MAPE), which quantifies prediction error by comparing the model’s expectation (Q-value) with the observed reward (BOLD response) for each trial (see Table 1); (ii) the convergence of the reinforcement learning algorithm (see Figure 4); and (iii) the Q-value trajectory for the final state-action pair (see Figure 5). Except for MAPE, we used the dashboard component to monitor these and other metrics (e.g., motion estimation, learning curve) in real-time during data acquisition (see Supplementary Figure 5).

**Figure 5:**
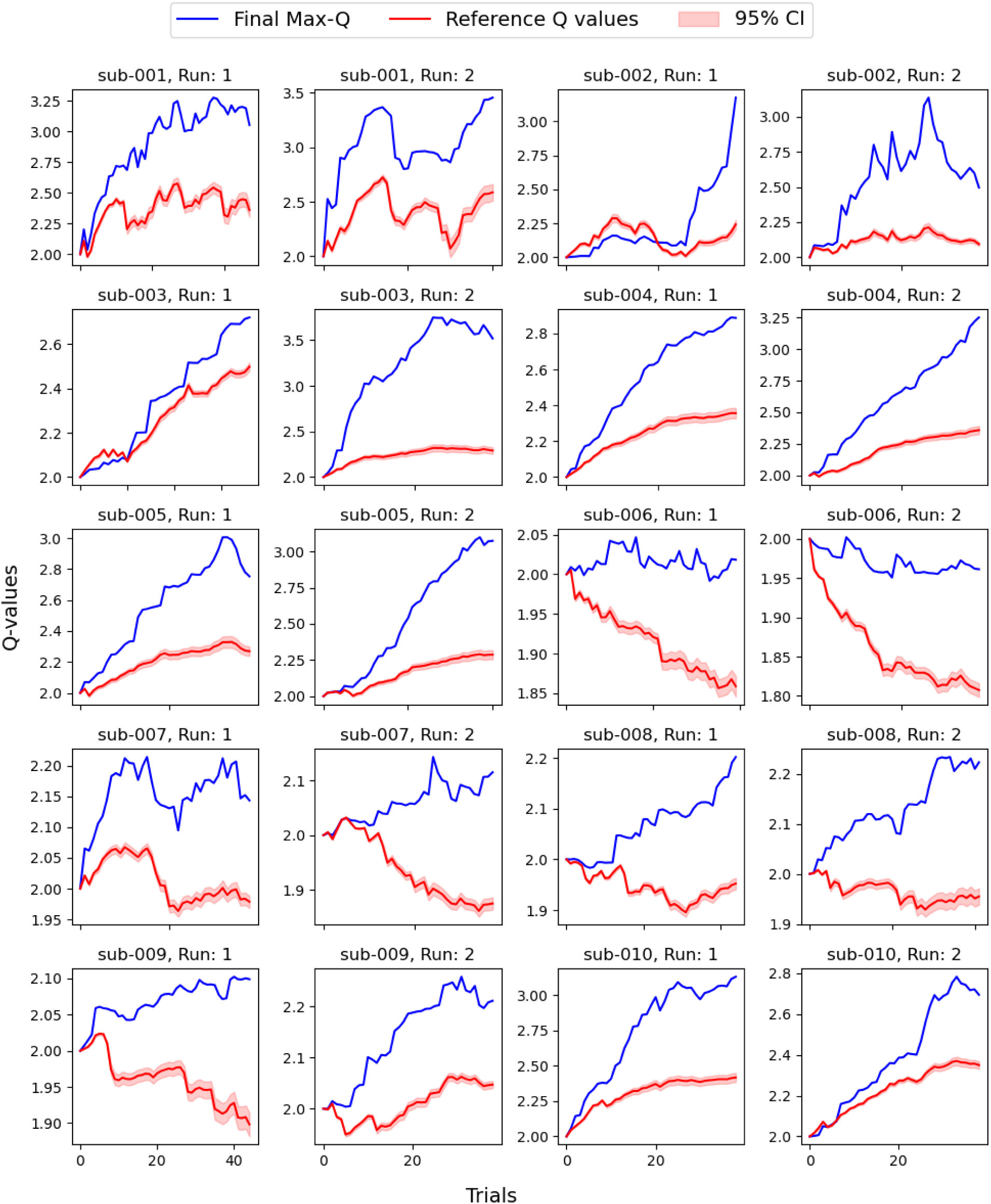
Evolution of Q-values during learning. The figure shows the Q-value corresponding to the location of the maximum Q-table value in the final trial (blue line) compared with the mean Q-value across all other stimulus configurations that do not correspond to the optimal region (i.e., excluding the maximum and its neighboring values; red line with 95% confidence intervals). The trajectories illustrate how, during learning, Q-values associated with the optimal stimulus region increase or remain stable, whereas Q-values for suboptimal configurations decrease or remain lower. This separation of value estimates enables the agent to progressively exploit stimulus configurations expected to maximize the target brain response while maintaining the learned structure of the stimulus space. This differentiation between optimal and non-optimal stimulus configurations was highly significant across all runs and all participants in the proof-of-concept study, suggesting that RLBF may provide a promising framework for individualized stimulus optimization based on participant-specific neural responses.

**Table 1:** Inter-trial Mean Absolute Percentage Error (MAPE) improvement and Signal-to-Noise Ratio (SNR) per run. MAPE improvement reflects the trend of error reduction across trials, calculated via permutation testing on MAPE slopes, yielding one-tailed p-values. While the learner generally converged near expected contrast values, noisy data occasionally hindered consistent improvement. The SNR column provides a proxy for empirical noise levels during fMRI acquisition, based on the ratio of mean reward to its standard deviation. Regardless of the SNR levels in the incoming data, the algorithm consistently identifies maximum contrast as shown by the position of the highest Q-table value at the last trial. For details about the calculations and code, see our supporting repository [34].

| Subject | Run | MAPE Slope (Improvement) | P-value | SNR | Final Position | Q-Table Max Position |
| --- | --- | --- | --- | --- | --- | --- |
| sub-001 | 1 | -0,0165 | 0,002 | 3,757 | [0,4, 0,5] | [0,9, 0,5] |
| sub-001 | 2 | -0,0090 | 0,003 | 3,394 | [0,7, 0,2] | [0,9, 0,0] |
| sub-002 | 1 | -0,0015 | 0,364 | 2,853 | [0,8, 0,8] | [0,9, 0,9] |
| sub-002 | 2 | -0,0013 | 0,312 | 3,486 | [0,8, 0,1] | [0,9, 0,0] |
| sub-003 | 1 | -0,0138 | 0,003 | 2,919 | [0,5, 0,1] | [0,9, 0,0] |
| sub-003 | 2 | -0,0072 | 0,002 | 4,631 | [0,9, 0,9] | [0,9, 0,9] |
| sub-004 | 1 | -0,0086 | 0,001 | 6,850 | [0,7, 0,4] | [0,9, 0,0] |
| sub-004 | 2 | -0,0095 | 0,001 | 4,947 | [0,9, 0,9] | [0,9, 0,9] |
| sub-005 | 1 | -0,0157 | 0,032 | 3,181 | [0,9, 0,8] | [0,9, 0,9] |
| sub-005 | 2 | -0,0050 | 0,013 | 4,152 | [0,8, 0,9] | [0,9, 0,9] |
| sub-006 | 1 | -0,0065 | 0,227 | 4,030 | [0,7, 0,3] | [0,9, 0,0] |
| sub-006 | 2 | -0,0169 | 0,003 | 4,280 | [0,8, 0,3] | [0,9, 0,9] |
| sub-007 | 1 | -0,0142 | 0,092 | 3,259 | [0,9, 0,2] | [0,9, 0,0] |
| sub-007 | 2 | 0,0085 | 0,708 | 3,138 | [0,1, 0,7] | [0,9, 0,9] |
| sub-008 | 1 | -0,0014 | 0,389 | 4,026 | [0,8, 0,6] | [0,9, 0,9] |
| sub-008 | 2 | -0,0037 | 0,243 | 4,092 | [0,7, 0,3] | [0,9, 0,9] |
| sub-009 | 1 | 0,0026 | 0,665 | 4,528 | [0,8, 0,2] | [0,9, 0,9] |
| sub-009 | 2 | -0,0071 | 0,082 | 4,065 | [0,7, 0,8] | [0,9, 0,9] |
| sub-010 | 1 | -0,0039 | 0,221 | 3,721 | [0,9, 0,2] | [0,9, 0,0] |
| sub-010 | 2 | 0,0002 | 0,528 | 4,208 | [0,9, 0,3] | [0,9, 0,0] |

As the reinforcement learning agent interacts with the stimulus space in order to maximize rewards, a decrease in average absolute percentage error is a sign of successful learning, both during the simulation and in real-time fMRI scanning. The prediction error, represented by the decrease in MAPE across trials, indicated that the algorithm effectively refined its actions to maximize brain activity in the primary visual cortex (V1). However, fluctuations in prediction error are expected even in the final trials, as the soft-Q learner, depending on the temperature parameter, may keep on exploring even with a well-learned Q-table.

Nevertheless, we have quantified the convergence of the reinforcement learning algorithm by calculating the Euclidean distance between the coordinates of consecutive actions in the Q-table space. Specifically, we check if the algorithm converged within a moving window, where the distance is averaged (window size=2). Stable convergence values of 0, indicating consistent selection of the same action, are considered as a sign of successful convergence of the algorithm. In the simulation, convergence reaches a stable value of 0 in the final trials, showing that the algorithm rapidly identifies potential optimal actions. For the empirical data acquired during real-time fMRI scanning, we applied a slightly more liberal convergence threshold, as with real data the model didn’t reach the level of confidence required to entirely stop exploring (mean convergence 0.25).

This slight deviation from zero was anticipated, given the inherently noisier nature of fMRI data compared to the simulated data. Interestingly, this discrepancy occurs despite the Signal-to-Noise Ratio (SNR) hyperparameter in the simulation being set to 3.30, lower than the average SNR of 4.02 found in the empirical data. Nonetheless, both the simulation and empirical results show consistency, demonstrating that the algorithm successfully converges within the targeted time frame of approximately 10 minutes of scanning in all participants. This consistency (shown in Figure 3 and by the position of the highest q-value at the last trial in Table 1) does not depend on the early states of the Q-table, as the whole process starts with the agent selecting a random action, and indicates generally stable performance, with the reinforcement learner typically identifying actions associated with maximizing contrast as optimal. This is reflected in the higher Q-values associated with high-contrast actions in the average Q-table. Furthermore, this trend towards optimizing contrast is observable within the optimization trajectories of individual participants (see Table 1; data available in our supporting repository [34]).

Although convergence serves as the standard stopping criterion of our approach, setting the time constraint on each fMRI run is a critical factor in our proof-of-concept study. We set a limited scanning time of approximately 10 minutes as we anticipated that convergence would vary among participants. This variability was expected because the balance between exploration and exploitation is not only a function of the temperature parameter but also of the size and robustness of the brain responses. During this time, the maximum Q-values across participants aligned with the expected increase in contrast towards the maximum (i.e. a contrast of 1.0), while the frequency values shifted towards medium to high flickering frequencies (mean: 18Hz, SD: 10Hz), as anticipated by the literature on checkerboard stimulation in V1 [24,32,33].

Given the importance of fast convergence in our approach, we also assessed whether the algorithm can reach good solutions before the 10 minutes time limit. We observed that in all runs, when considering the position of the highest Q-value in the Q-table, the algorithm was able to identify both high contrast actions and the medium-high frequency range (mean: 19Hz, SD: 9Hz) after approximately 15-25 trials, or around 4-5 minutes of scanning time (see Supplementary Figure 4). This rapid progression to the optimal solution is a key feature of our approach, as it demonstrates, at least within the simple setting of our proof-of-concept study, that it is possible to optimize the stimuli to drive neural activity in a specific direction at the level of a single participant, without the need for extended training times (See Figure 5). This represents an important feature for many potential applications [8] and whether it is possible to maintain low scanning times has to be confirmed in future studies involving more complex stimulus spaces.

## 6. Discussion

In this manuscript, we presented Reinforcement Learning via Brain Feedback (RLBF), an approach applying reinforcement learning to optimize experimental stimuli using real-time fMRI. This method leverages brain activity to dynamically adjust stimulus characteristics, proposing a first step towards AI-based stimulus optimizations that differ from traditional fixed-task paradigms where stimuli are predetermined. An important feature of this implementation is retraining the reinforcement learning agent for each new run, allowing the stimulus optimization process to potentially adapt to individual participant responses. While our proof-of-concept operates within a constrained, two-dimensional space targeting V1, the underlying principle, and related software implementation, aims to eventually enable navigation within more complex stimulus-spaces. The potential future integration with dynamic or interactive stimulation environments (e.g. videogames) or AI generated content, could theoretically open avenues for exploring novel stimulus configurations or real-time cognitive optimization, although demonstrating the efficacy and scalability of RLBF in such high-dimensional, less constrained scenarios is a goal for future research. In our proof-of-concept study, we use real-time BOLD signals from the primary visual cortex (V1) to optimize a flickering checkerboard stimulus. The reinforcement learning algorithm adjusts the contrast and frequency of the stimulus iteratively, thereby maximizing V1 activity. This optimization occurs within a relatively short timeframe, with the algorithm identifying high-contrast actions within approximately the 10-minute time limit. Notably, the results from our empirical measurements align with existing knowledge on the contrast and frequency dependence of the checkerboard response [24,32,33], providing initial confirmation of the feasibility of our proposed approach. Nevertheless, while RLBF yields promising results in its current configuration, the relatively simple stimulus space used in this proof-of-concept study does not fully capture the complexity of more challenging applications. This limitation becomes particularly relevant when considering high-dimensional stimulus spaces with continuously varying parameters, where optimization may be further complicated by higher-order cognitive factors, such as the influence of the brain’s own predictive processes on the measured neural response. The extent to which RLBF can scale to such complex stimulus spaces in single-participant real-time fMRI settings remains an important question for future research.

Our extendable and highly portable software implementation and the presented proof-of-concept study paves the way for future efforts. While in the present proof-of-concept study we deliberately focused on approximating simple solutions, the statistical power necessary for future efforts with more complex stimulus spaces may be achieved by improved fMRI processing and reinforcement learning approaches and, importantly, by multi-region or network-level brain features (e.g. multivariate brain signatures [37] or computational model-based readouts, like functional attractors [38,39]). Building on our current findings, the next step is to explore cognitive modulation during exposure to dynamic tasks or AI-generated content within the RLBF framework, which could greatly expand its applications in both basic and translational research.

In basic research, RLBF could provide a novel approach for systematically probing the relationship between stimulus features and neural representations in adaptive, closed-loop settings. Rather than relying exclusively on predefined paradigms, RLBF may help characterize the stimulus selectivity, specificity, and contextual dependence of neural responses. For example, it could be used to investigate whether regions traditionally associated with specific functions, such as the fusiform cortex in face perception [40] or the amygdala in threat processing [41], exhibit broader or more context-dependent response profiles when explored across richer and less constrained stimulus spaces. Similarly, RLBF could provide a framework for evaluating the specificity and robustness of multivariate neural signatures (like 42 and 43) by identifying stimuli that maximize, minimize, or dissociate predictive readouts from their intended psychological constructs. This could be particularly valuable for investigating the generalizability and potential vulnerabilities of brain-based biomarkers [44,45,46,47], including the possibility of identifying adversarial or out-of-distribution stimulus configurations that challenge signatures such as pain or affective predictive models.

Beyond basic characterization of neural representations, RLBF may offer new opportunities for studying adaptive behavior and learning in complex environments. By allowing stimulus spaces to be optimized while individuals interact with dynamic environments, RLBF could help investigate how neural systems adapt during decision-making under uncertainty, skill acquisition, and cognitive control [48,49,50]. For example, it may contribute to the discovery of task features that promote optimal performance in naturalistic environments, including interactive games or complex sensorimotor tasks, or help identify and maintain neural states associated with efficient learning and adaptation. Similarly, in brain-computer interface applications, RLBF could potentially assist in identifying stimulus configurations that maximize the discriminability or reliability of neural signals used for communication or control.

In translational research, RLBF may enable individualized characterization and optimization of brain-stimulus relationships. Potential applications include identifying personally relevant fear or anxiety-provoking stimulus features [51,52] that could inform subsequent exposure-based interventions, or mapping individualized sensitivity profiles that may improve the targeting of therapeutic strategies. RLBF could also be combined with generative AI-based stimulus spaces to create personalized sensory environments, for example, optimizing music or visual experiences to support analgesia, relaxation, emotional regulation, or cognitive engagement [53]. Other potential applications include adaptive calibration of neurotechnology systems, individualized optimization of neurofeedback protocols, and the development of personalized cognitive training approaches [54].

The examples discussed above are intended as illustrative possibilities that motivate future research directions rather than immediate applications. Their feasibility, specificity, scalability, and clinical utility remain open questions that will require systematic validation in future studies. Nevertheless, the concept of RLBF introduces a conceptual shift from treating stimulus design as a fixed experimental choice toward viewing it as an adaptive optimization problem, providing a framework for exploring individual and population-level brain-stimulus relationships in a more systematic and flexible manner.

## 7. Conclusion

In this work, we introduced the RLBF concept, developed a modular and extensible software implementation, and demonstrated its feasibility in a proof-of-concept study using real-time fMRI feedback to guide stimulus optimization within a single scanning session. While the present study intentionally focused on a simple stimulus space, these results provide an initial foundation for exploring the promises and limitations of adaptive, neural-driven experimental design in more complex settings. By enabling more targeted and flexible interrogation of brain responses, RLBF may help bridge the gap between controlled laboratory paradigms and more ecologically valid, individualized applications, while offering new opportunities to evaluate the specificity of neural representations and biomarkers. Combined with ongoing advances in artificial intelligence and generative modeling, RLBF may further contribute to the development of personalized approaches for studying and modulating brain function. The scalability, robustness, and practical utility of these applications remain important topics for future research.

## Ethics

This study was conducted in accordance with the ethical principles outlined in the Declaration of Helsinki. Ethical approval for the research was obtained from the Ethical Committee of the University of Duisburg-Essen. Informed consent was obtained from all participants prior to their involvement in the study.

## Data accessibility

All materials, code and data are available on GitHub, including the main RLBF software repository (git.uni-due.de/pnl-lab/rlbf) and the repository containing the analyses presented in the manuscript (git.uni-due.de/ht3489/RLBF_analysis).

