## Supplementary material for "Reinforcement Learning via Brain Feedback for real-time fMRI-based adaptive stimulus generation"

### Supplementary Materials

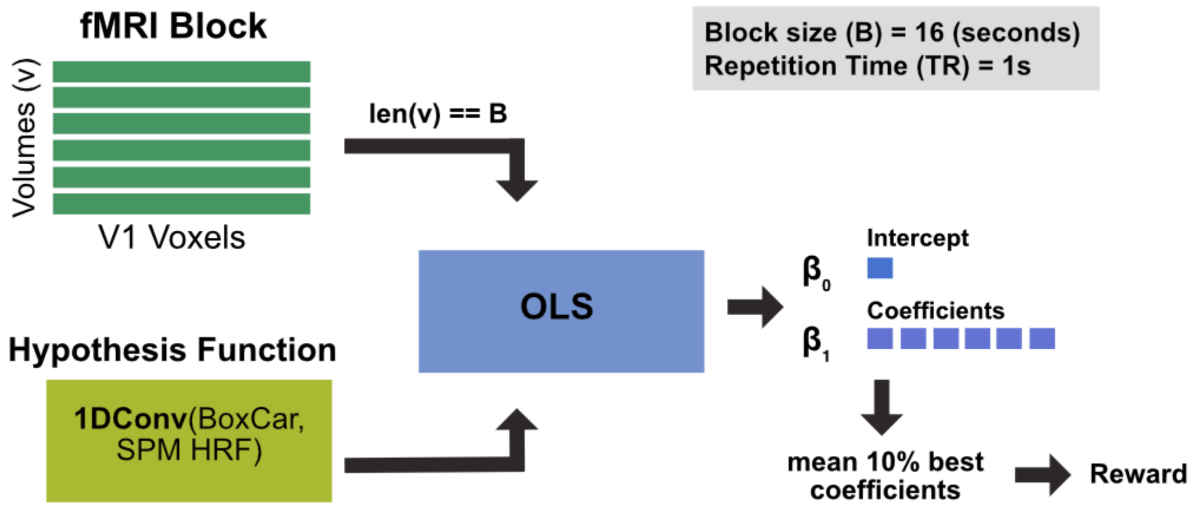


**Supplementary Fig.1:** Diagram showing reward calculation. A hypothesis function, representing the expected BOLD response (derived from convolving the experimental block design with the canonical SPM hemodynamic response function), is defined before the real-time process starts. During acquisition, primary visual cortex (V1) fMRI data for each volume is fitted to this function using an Ordinary Least Squares (OLS) model. The reward signal is calculated by averaging the coefficients corresponding to the top 10% best fit (see the 'real-time data processing protocol' section in the manuscript).

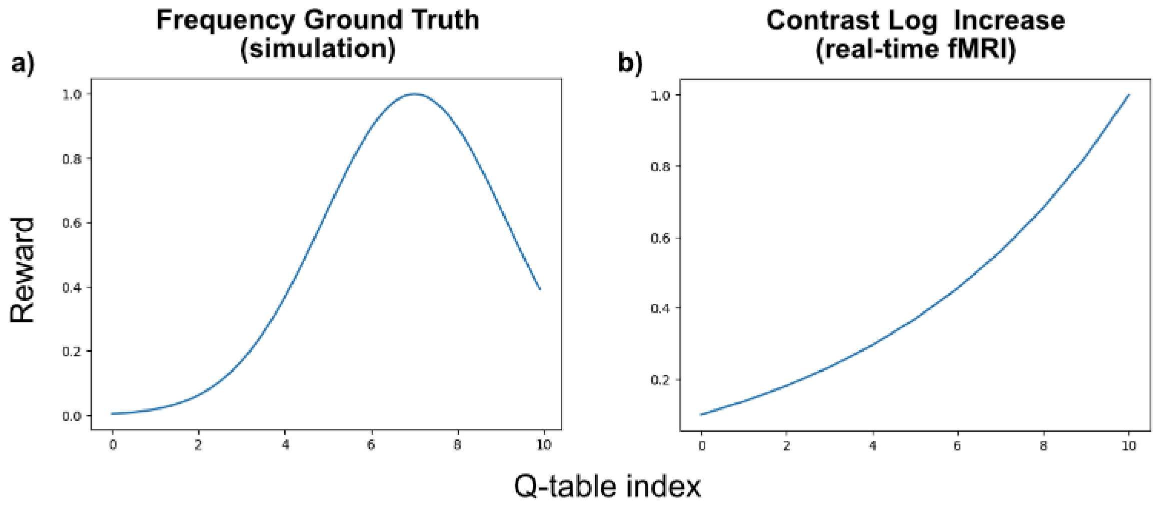


**Supplementary Fig.2:** The non-linear ground truth for frequency tuning used in simulations, where the reward peaks at an arbitrary value of 0.7 (**a**). Logarithmic scaling of checkerboard contrast presented during the real-time fMRI experiment (**b**), mimicking the visual system's non-linear Contrast Response Function (CRF) [1].

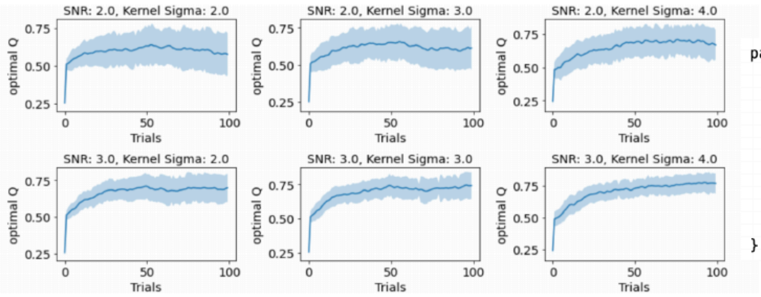


```
parameters = {
    'snr': [0.5, 1, 2, 3, 5],
    'n_bins': [10],
    'kernel_size': [20],
    'kernel_sigma': [0.5, 1.0, 2.0, 3.0, 4.0],
    'learning_rate': [0.05, 0.1, 0.3, 0.4, 0.8, 0.9],
    'temperature': [0.2],
    'min_temperature': [1e-5],
    'max_temperature': [1.0],
    'reduce_temperature': [False],
    'decay_rate': [0.001]
}
```

**Supplementary Fig.3:** Simulation outcomes (100 trials) showing Q-values under the optimal condition (frequency=0.7, contrast=1.0) and with optimal hyperparameters. Higher resulting Signal-to-Noise Ratio (SNR > 2) and Smoothing σ (kernel sigma) indicates better model performance. Key tuned hyperparameters included: input SNR (controlling simulated noise level), kernel sigma (determining Q-table smoothing extent), and learning rate (governing Q-value update step size). Other parameters like temperature and size of the smoothing kernel (kernel size) were fixed, while temperature decay and decay rate were unused in this study (For all the results of our hyperparameter tuning see [2]).

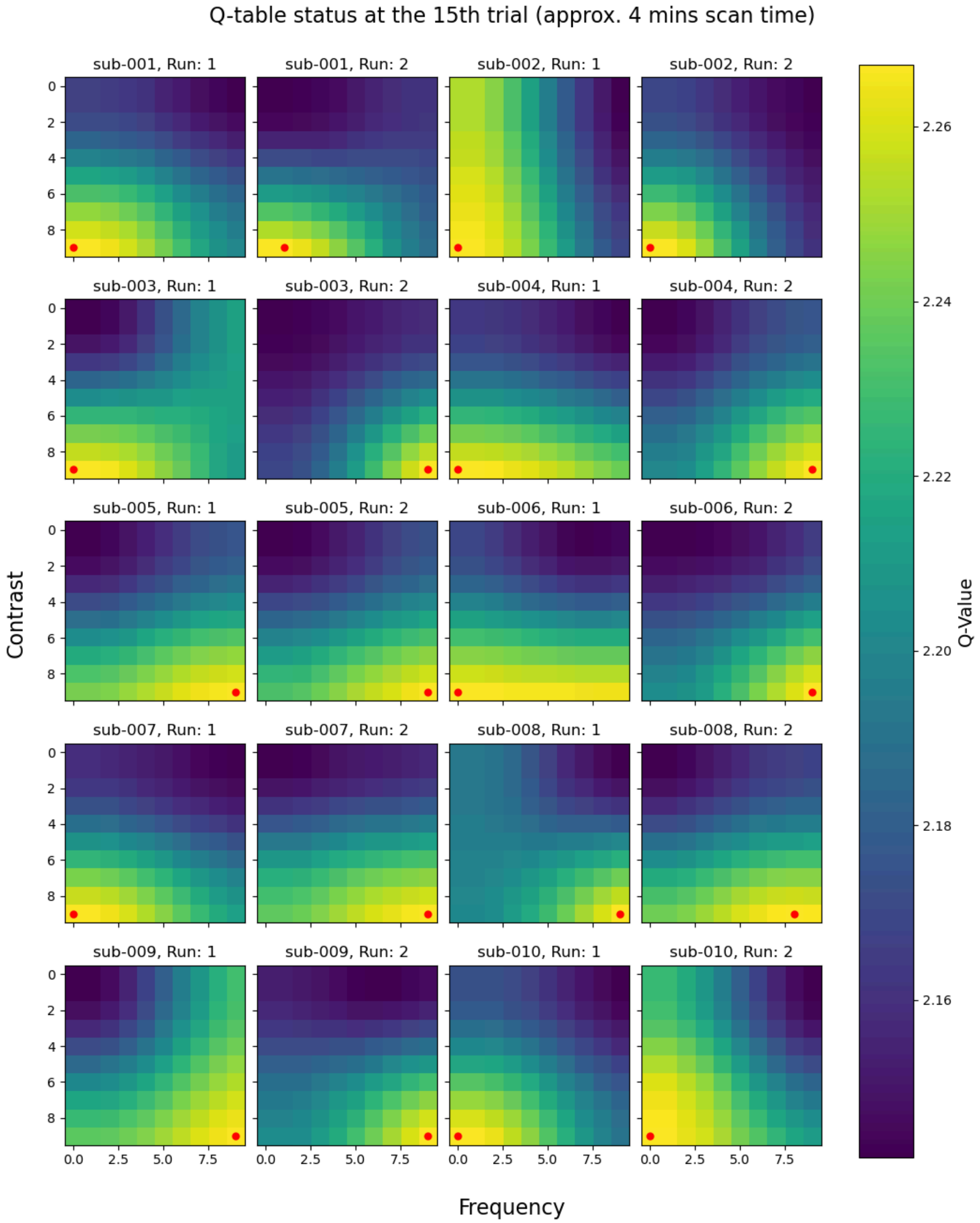


**Supplementary Fig.4:** illustrates the status of the Q-table for each participant and run at the 15th trial (approximately 4 mins scanning time). The colors in the Q-table reflect the Q-values, with brighter colors indicating higher values, which the RL agent considers more likely to cause a positive change in brain response. The red dot indicates the position of the maximum Q-value at the current state, showing the action with the highest softmax probability of being selected as the next action. The figure shows that the Soft-Q-learner can quickly identify the range of optimal choices, around high-contrast, on the single subject and in a very limited amount of scanning time.

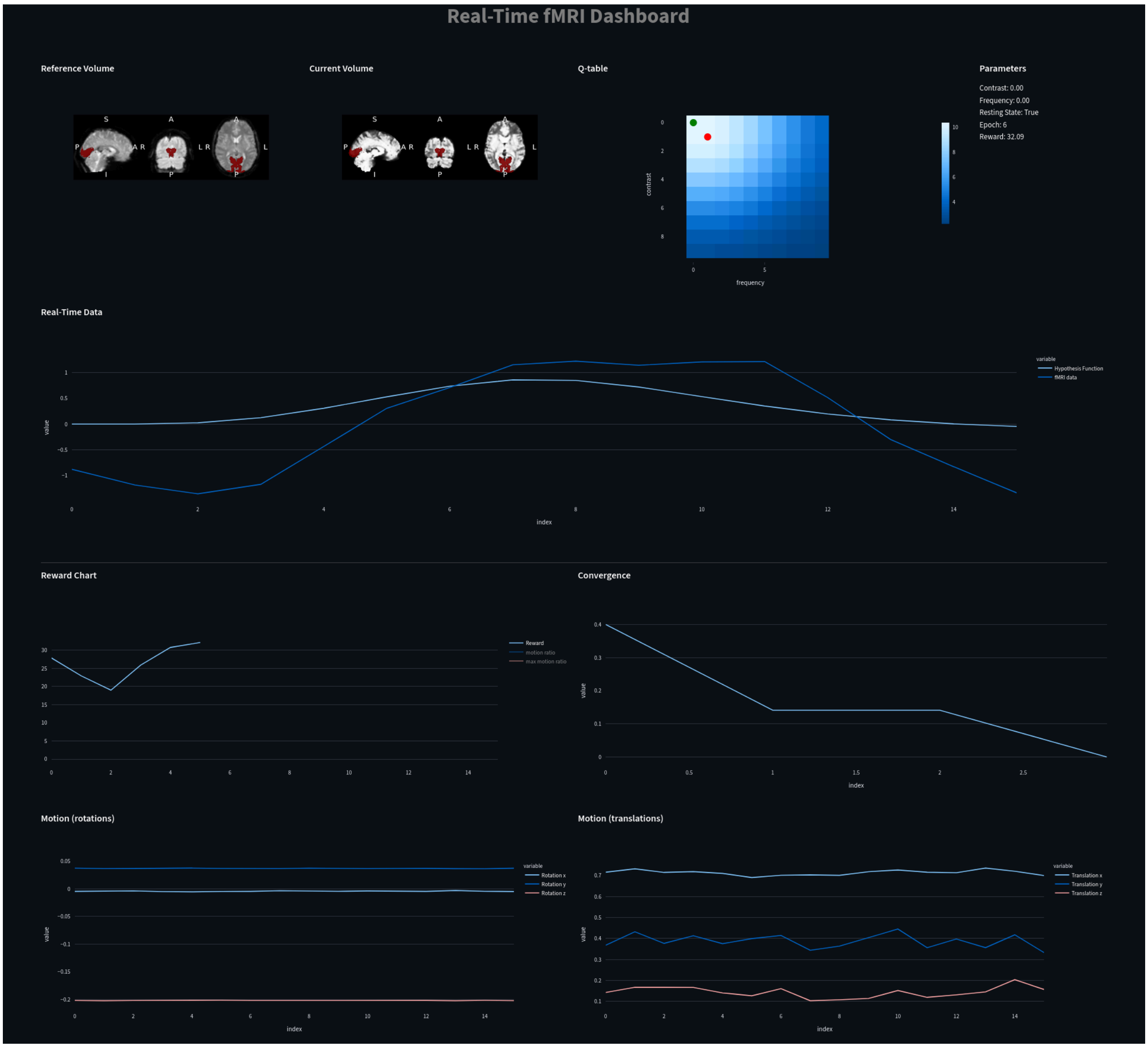


**Supplementary Fig.5:** Preview of the **real-time fMRI dashboard** specifically showcasing part of the interface utilized during the proof-of-concept study for real-time data acquisition. The dashboard displays (**top**) the functional reference volume (registered to MNI152), a real-time preview of the current, rank-harmonized, volume (updated every TR=1s), the Q-table showing the Agent's walk through the stimulus space, and updated study parameters (e.g., contrast, frequency, epoch). It also presents the timeseries for a block of fMRI data (16s total: 11s rest, 5s stimulation) overlayed on the hypothesis function, standardized for visualization and other plots, reporting the overall progress of the agent (i.e. reward and convergence) and motion regressors (**bottom**). Developed using Streamlit [3], this dashboard is highly customizable and allows users to embed their preferred data visualization.

2. Gallitto G, Englert R, Kincses B, Kotikalapudi R, Li J, Hoffschlag K, Ali S, Bingel U, Spisak T. Supporting repository for the manuscript "Towards generative AI-based fMRI paradigms: reinforcement learning via real-time brain feedback" [Software]. Retrieved from https://git.uni-due.de/ht3489/RLBF_analysis.

3. Streamlit [Software]. Retrieved from https://streamlit.io/. Accessed Jan 11, 2024.
